# Computational Correction of Spatially-Variant Optical Aberrations in 3D Single Molecule Localization Microscopy

**DOI:** 10.1101/504712

**Authors:** Ting Yan, Charles J. Richardson, Mingxing Zhang, Andreas Gahlmann

## Abstract

3D single-molecule localization microscopy relies on fitting the shape of point-spread-functions (PSFs) recorded on a wide-field detector. However, optical aberrations distort those shapes, which compromise the accuracy and precision of single-molecule localization microscopy. Here we employ a computational phase retrieval based on a vectorial PSF model to quantify the spatially-variance of optical aberrations in a two-channel ultrawide-field single-molecule localization microscope. The use of a spatially-variant PSF model enables accurate and precise emitter localization in *x*, *y*- and *z*-directions throughout the entire field-of-view.

**OCIS Codes:** (100.6640) Superresolution; (180.2520) Fluorescence microscopy; (180.6900) Three-dimensional microscopy; (100.5070) Phase retrieval; (150.1135) Algorithms; (220.1010) Aberrations (global).

## 1. INTRODUCTION

Super-resolution microscopy represents a family of techniques employed to surpass the diffraction-limited spatial resolution of light microscopy through optical or computational means [1–3]. Single-molecule localization microscopy (SMLM) is one such technique, which has developed into a powerful tool in molecular and cellular biology [4–6]. In camera-based single-molecule localization microscopy, molecular positions are usually estimated by fitting a theoretical model to the experimentally-measured point-spread-functions (PSFs) of single fluorescent emitters. Determination of molecular positions with a precision of tens of nanometers enables the sub-cellular localization and co-localization of fluorescently-labeled biomolecules, as well as single-molecule tracking [7]. The accuracy and precision of molecular position estimation is of prime importance, because these metrics affect all subsequent steps in data analysis, such as estimation of cluster formation in pointillistic single-molecule localizations [8, 9], or molecular displacement estimation in single-molecule tracking [10–13]. The accuracy of the estimated position, i.e. its systematic deviation from the true position, is primarily dependent on the choice of the theoretical PSF model used for fitting the measured fluorescence intensity data [14]. The precision of the estimated localization, i.e. its random statistical variability upon repeated measurements, is primarily determined by the signal-to-background ratio of the fluorescence intensity data [4–6]. Efforts to improve localization accuracy and precision require minimizing systematic and random errors during data acquisition and computational post-processing [15].

A major advancement of single-molecule localization microscopy has been the ability to use PSF engineering to encode the 3D position of an emitter and/or its emission wavelength [16–18] or its molecular orientation [19–21]. To achieve emitter localization below the diffraction limit in both the lateral (*x*,*y*) and axial (*z*) dimensions, PSFs have been engineered to change shape rapidly along the z-dimension, either through stretching or through rotation [22–30]. For example, the double-helix point-spread-function (DHPSF) was engineered to produce two high-intensity lobes that revolve around the emitter’s *x*,*y*-position in a *z*-displacement dependent manner [24, 25]. Careful calibration measurements have revealed that the fitted *x,y*-position, obtained using analytical PSF models, deviates slightly from the true lateral position of an emitter as a function of *z*-displacement. This ‘wobbling’ behavior of double-helix and astigmatic PSFs can be calibrated and corrected to obtain more accurate estimates of an emitter’s lateral position [31, 32].

The large chip size of modern sCMOS camera has enabled SMLM in ultrawide fields-of-view (diameters of up to 250 µm) [33–35]. Ultrawide-field imaging combined with fast frame rates (up to 100 Hz for full chip readouts), has helped increase the overall data acquision throughput [36]. However, the high-NA objective lenses used for SMLM cannot offer full aberration correction everywhere in the field-of-view. It is well established that optical aberrations become more pronounced as emitters are localized far away from the central optical axis [33, 37–40]. Thus, the prevalent assumption of a spatially-invariant PSF in the imaging system does not hold and the PSFs used in high-precision, high-accuracy SMLM need to be treated as spatially-variant.

Previous work extracted emitter positions from experimental PSFs using simple analytical, interpolated, or scalar/vectorial PSF models. Simple analytical models typically use Gaussian functions to approximate the experimentally measured fluorescence intensity profiles [4–6]. More sophisticated models based on the Gibson-Lanni model [41] or Hermite functions [42] have also been employed. Simple analytical functions are however not flexible enough to accurately model the complex shapes of aberrated PSFs encountered in real microscopes. To overcome these limitations, piecewise polynomial functions have been used to interpolate experimentally-acquired PSFs [43–46]. Such interpolated model functions contain larger numbers of adjustable parameters compared to the simple analytical functions, which makes them more flexible to approximate the PSF intensity profiles. Finally, models based on scalar or vectorial diffraction theory explicitly account for the propagation of light through high numerical aperture imaging systems [16, 18, 47]. Augmented vectorial models also include the glass-water refractive index boundary at the microscope coverslip to enable estimation of PSF shapes for emitters located within an index mismatched medium [48, 49]. However, neither interpolated or diffraction-theory-based models have addressed the spatial variance of experimental PSFs in SMLM.

Here, we use a vectorial PSF model and refine it using computational phase retrieval to quantify the spatially-variant optical aberrations in two different color channels of a single-molecule localization microscope. We show that the spatially-variant vectorial PSF model enables accurate and precise 3D emitter localization throughout an ultrawide field-of-view. We further show that the same emitters can be co-localized in two different color channels with an accuracy less than 25 nm, which matches localization precisions typically achieved in live-cell SMLM. Our work provides a quantitative framework for processing multi-channel 3D SMLM data acquired with arbitrarily engineered PSFs in an ultrawide field-of-view.

## 2. THEORY AND METHODS

### 2.1 Optical Setup

Experiments were performed on a custom-built dual-color inverted fluorescence microscope based on the RM21 platform (Mad City Labs, Inc) [Fig. 1]. Immersion oil was placed between the objective lens (UPLSAPO 60X 1.4 NA, Olympus) and the glass cover slip (#1.5, 22 mm x 22 mm, Schott). The sample was excited with a 514 nm laser (Genesis MX514 MTM, Coherent) and a 561nm laser (Genesis MX561 MTM, Coherent). All excitation lasers were circularly polarized by zero order quarter wave plates (WPQ05M-514 and WPQ05M-561, Thorlabs), and a bandpass filter filtered the spectral profile of the 514 nm laser (ET510/10bp, Chroma).

**Fig. 1.**
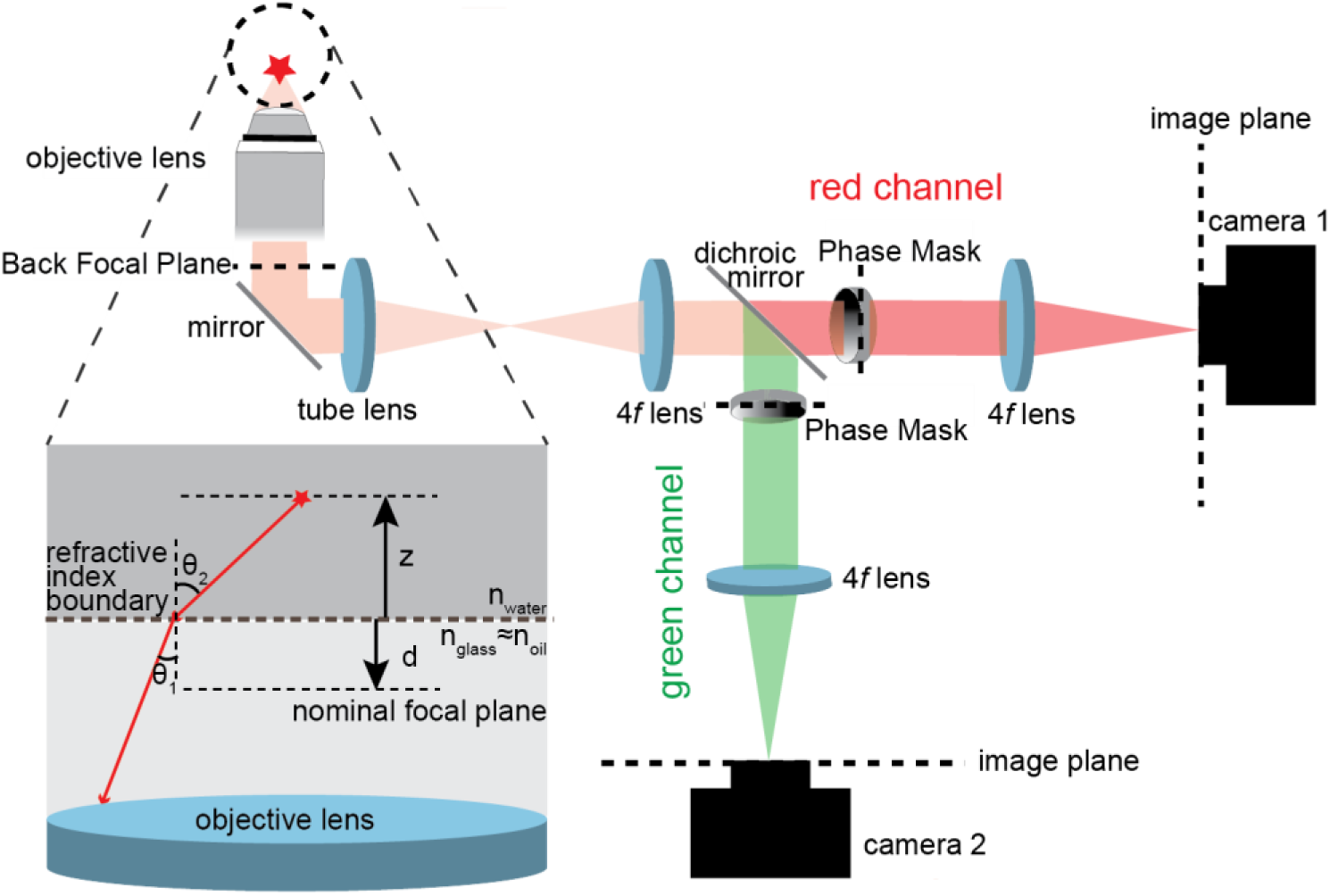
Schematic fluorescence emission paths of the microscope system. Two phase masks are placed in planes conjugate to the back focal plane of the objective lens (dashed lines) to generate the DHPSF. The inset shows the relative distance *d* of nominal focal plane relative to the refractive index boundary, as well as the emitter depth z in the refractive index mismatched medium. (The nominal focal plane is equal to the focal plane in absence of a refractive index boundary.) Positive values of *d* place the nominal focal plane below the refractive index boundary, whereas a negative distance *d* places the nominal focal plane into the refractive index mismatched medium. Emitter depths are parameterized by positive values of *z*, i.e. they are always localized above the refractive index boundary.

Fluorescence emission was passed through a shared filter set of LP02-514RU-25 (Semrock), NF03-561E-25 (Semrock), and ET700SP-2P8 (Chroma). A dichroic beam splitter (T560lpxr-uf3, Chroma) was then used to split the emission pathway into ‘red’ and ‘green’ channels, corresponding to the transmitted and reflected light, respectively. The ‘red’ transmitted channel contained an additional 561nm notch filter (ZET561NF, Chroma) to block scattered laser light. Each emission path contains a wavelength-specific dielectric phase mask (Double Helix, LLC) placed in the Fourier plane of the microscope and mounted on high-precision XYZ translation stages (9062-XYZ, Newport) to generate a double-helix point-spread-function [25, 50]. The fluorescence signals in both channels are detected on two separate sCMOS cameras (ORCA-Flash 4.0 V2, Hamamatsu).

### 2.2 Sample Preparation and Imaging

For samples consisting of calibration beads, we diluted 100 nm tetraspeck fluorospheres (Invitrogen) diluted in PBS at a ratio of 1:3000 (v/v) and then spin-coated them onto a microscope coverglass. To maintain a refractive index similar to water, we covered the beads with a 1.5% (w/v) low-melting point agarose (Fisher Scientific) pad made with PBS buffer to mimic experimental conditions used for live cell imaging [12, 51, 52]. To generate a sample of beads immobilized away from the microscope coverslip, we suspended fluorescent beads into molten 1.5% agarose and spotted a 10 µl of mixture onto the coverglass which was previously spin-coated with fluorescent beads.

The phase mask lateral position was optimized prior to each experiment by scanning the nominal focal through plane a single fluorescent bead immobilized on the microscopy coverlip. The resulting PSF sections were analyzed using the Easy-DHPSF software [53]. The phase masks lateral position was adjusted to ensure even lobe intensities and minimal lateral shifts of the fitted positions at different positions of the nominal focal plane.

Calibration images to determine the spatially-variant aberrations profile were acquired by scanning the nominal focal plane through fields-of-view containing many immobilized fluorescent beads. Specifically, we stepped the distance *d* in 50 nm increments through a total range of 3 µm and recorded 10 PSF images at each step. All images were acquired with an exposure times of 30 ms and illumination intensities of 0.5-4.5×10^6^ mW/cm^2^.

### 2.3 Modelling an Isotropic Emitter

We employ a vectorial model of light propagation to simulate the PSF of an emitter in a refractive index mismatched medium [48, 49]. Briefly, we simulate polarized emission of a dipole emitter and propagate the light through the refractive index boundary. The refracted light waves are then collected by a high numerical aperture (NA) objective lens. This vectorial PSF model provides the amplitudes and phases of the light waves in the back focal plane (BFP) of the objective lens, i.e. the pupil function [Fig. 1]. As detailed further below, the phases of these waves are dependent on the position of the emitter, whereas the amplitudes are not. The light waves are then further propagated through the imaging system (one tube lens and two 4f lenses) using Fourier optics to ultimately yield the final intensity profile in the image plane. To model the PSF of an isotropic emitter, the PSFs of three specific dipole orientations are summed in the image plane [48, 54]. We use this model to compute the PSFs of isotropic emitters at arbitrary *x*,*y*,*z*-positions above the refractive index boundary [Fig. 1 inset]. The mathematical detail of the model are described in the following.

The refraction and transmission of the emitted light at the refractive index boundary is described, respectively, by Snell’s law, *n*_1_ *sinθ*_1_ = *n*_2_ *sinθ*_2_, and the Fresnel coefficients. The Fresnel coefficients for transmission of *s*- and *p*-polarized light are given by

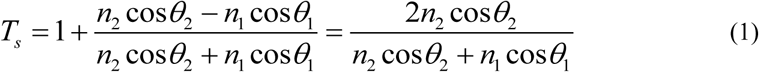

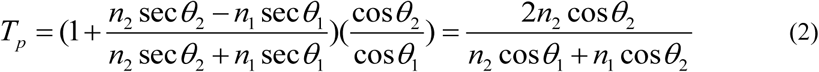

where *n*_1_ = *n*_*glass*_ ≈ *n*_*oil*_ = 1.52, *n*_2_ = *n*_*water*_ = 1.33. The variables *θ* and *ϕ* correspond to the polar and azimuthal angles of emitted light rays [Fig. 1(a)]. The polar angle *θ* maps to the BFP polar coordinate *ρ* through *ρ* = sin*θ*_1_ = *n*_2_ sin*θ*_2_ / *n*_1_, whereas the azimuthal angle *ϕ* maps one-to-one to the BFP azimuthal coordinate. The numerical aperture of the objective lens defines the range of angle *θ*_1_ that can be collected and thus the maximum radius *ρ*_*max*_ in the BFP according to

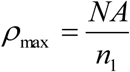

The electric field amplitudes of *x* and *y* polarized light, 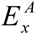 and 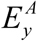, are calculated using,

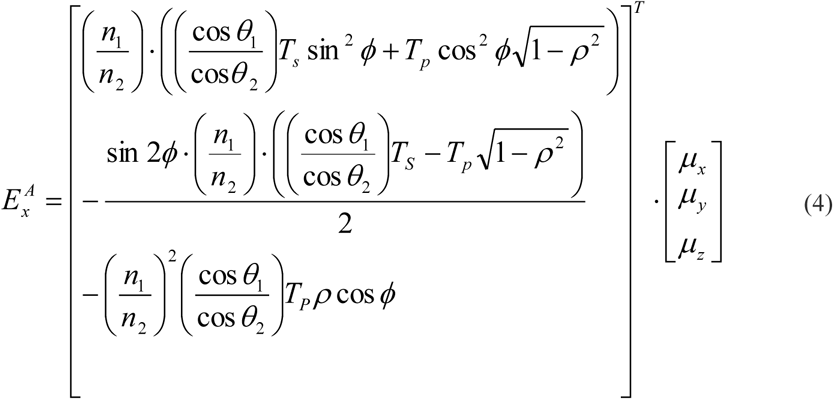

and

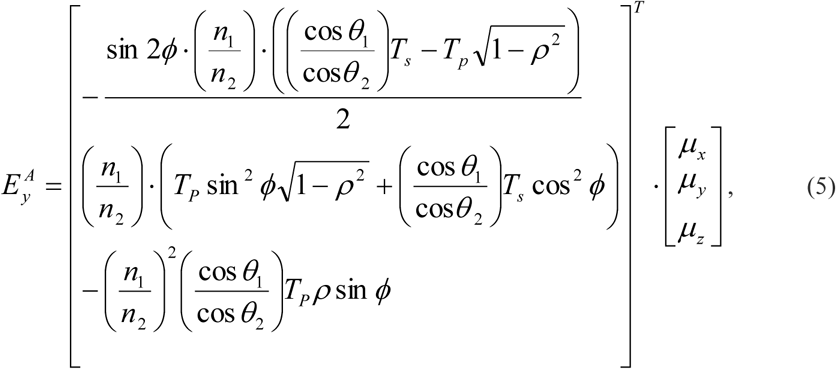

respectively, where *µ*_*x*_, *µ*_*y*_, and *µ*_*z*_ represents the emitter’s dipole orientation in Cartesian coordinates. Alternatively, the dipole orientation can also be defined in spherical coordinates according to

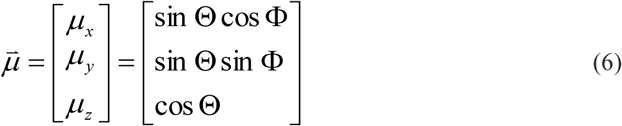

The phases of the electric field depend on several factors, namely the *z*-position of the emitter above the refractive index boundary

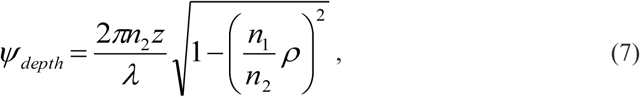

and the distance *d* between the nominal focal plane and the refractive index boundary

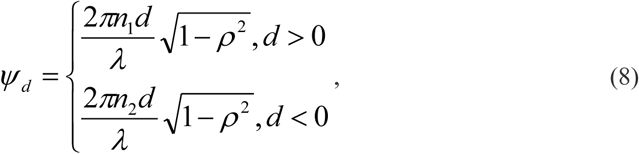

and the lateral *x*,*y*-displacement of the emitter from the central optical axis

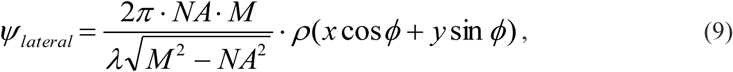

where in these expressions *λ* is the emission wavelength and *M* is the magnification of the imaging system.

The overall phase in the BFP is thus given by

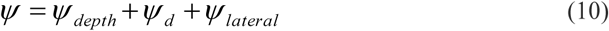

And the complex amplitudes in the BFP are then given by

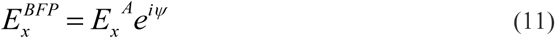

and

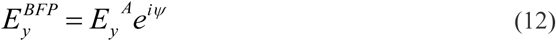

To compute the PSF in the image plane, we propagate the light wave through the rest of the imaging system using Fourier optics

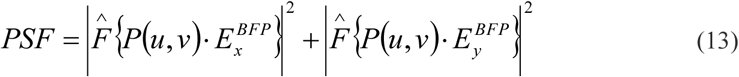

where *P(u,v)* specifies the phase manipulation, which, in our case, alters the standard PSF to the DHPSF.

The model described so far produces PSFs assuming no optical aberrations.

To account for optical aberrations, Eqn 12 is modified to

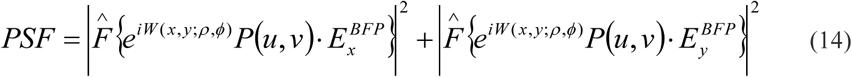

where *W*(*x*, *y*; *ρ*,*φ*) is the wavefront aberration phase which depends on both specimen plane and back focal plane coordinates, described by

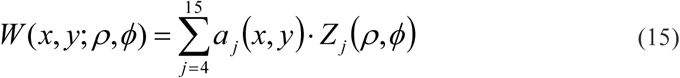

here for each Noll index (*j*), *a*_*j*_(*x,y*) is the spatially-variant aberration coefficient, and Z_*j*_ is the Zernike polynomial normalized to π.

In the vectorial PSF model, we assume that the fluorescence emission from each emitter in our case is isotropic. Isotropic emitters are modelled by summing the PSFs of three anisotropic emission dipoles with orientations angle Φ and Θ of (0, π/2), (π/2, π/2), and (π/2, 0) (See Eqn 6). Additionally, a Gaussian blur with radius of 0.58 pixels (62.6 nm) was added to the simulated image to account for the finite spherical volume of fluorescent beads. This blur radius was determined by minimizing the difference between a simulated 100 nm sphere filled with isotropic emitters and a blurred isotropic point emitter in the absence of noise and aberration.

### 2.4. Data Analysis

We used two different methods to fit the DHPSFs in order to extract the *x*,*y*,*z*-positions of individual emitters. The first method is the analytical model built into the Easy-DHPSF fitting routine, namely a model consisting of two laterally displaced Gaussian functions [53]. This routine was modified so that both least squares (LSQ) and/or maximum likelihood estimation (MLE) can be performed [36]. The other method uses the spatially-variant vectorial PSF model described above and MLE for PSF fitting. To determine the *x,y,z*-positions of individual emitters accurately using the vectorial PSF model, the distance *d* of current frame and the aberration coefficients *a*_*j*_(*x,y*) (*j* = 4, … 15) of each emitter must be determined as described in the following.

## 3. RESULTS AND DISCUSSON

### 3.1 Quantifying Spatially-Dependent Aberrations in Large Fields-of-View

To determine the spatial dependence of optical aberrations in a large field of view, we immobilized fluorescent beads on glass coverslips and scanned the nominal focal plane of the objective lens over the beads in equally-sized (50 nm) steps. This procedure produced a set of 120 averaged images, each acquired at different defocus throughout the microscopes’ field-of-view. We then fit a 2-dimensional Gaussian function to the rotation angles of varying DHPSFs in each image and defined the apex of this Gaussian function as the position of the optical axis. The bead closest to the optical axis, which is assumed to be the least aberrated, was then used to estimate the nominal focal plane position *d* in each image, as well as the phase mask’s rotation and lateral displacement from the optical axis. MLE was used to minimize the difference between the experimental PSFs and unaberrated vectorial PSF model. To estimate the position of the nominal focal plane relative to the refractive index boundary, the *z*-position relative to the refractive index boundary) of the bead was assumed to be 50 nm (the diameter of the beads in our calibration measurements is 100 ± 6 nm, according to the manufacturer). The centermost bead’s *x*- and *y*-postions were also used as fitting parameters in this step.

Next, we quantified optical aberrations by estimating the coefficients *a*_*j*_(*x,y*) of the Zernike polynomials with *j* = 2,…,15 (Noll indices) at all bead locations by minimizing the difference between experimental PSFs and simulated PSFs, again using MLE [Fig. 2A], as described previously [18, 47]. After initial fitting, we did not consider the Zernike polynomials with Noll indices 2 and 3 further, because they correspond to tip and tilt, which do not change the shape of the PSF. To estimate the Zernike coefficients *a*_*j*_(*x,y*) with *j* = 4,…,15 at any location in the field-of-view, we fit a two-dimensional polynomial function of low order (usually 2-4, not to be confused with the Zernike polynomials) to the experimentally-determined coefficients [Fig. 2B] [55]. The order of polynomial surfaces was increased if necessary to minimize any spatial dependence of residuals [Fig. 2CD]. The distributions of residuals was analyzed to identify outlier beads with coefficients outside the mean ± 3 standard deviations range. Such beads were iteratively removed from the fit until all remaining beads fell within the above range. Excluded beads were removed from the surface fits of all other Zernike polynomials as well. The final polynomial surfaces can then each be queried at any location of the field-of-view to obtain the local aberration coefficients. The combined aberration coefficients provide the overall spatially-dependent wavefront aberration phase *W*(*x*, *y*; *ρ*,*φ*), which allows us to compute spatially-variant aberrated PSFs.

**Fig. 2.**
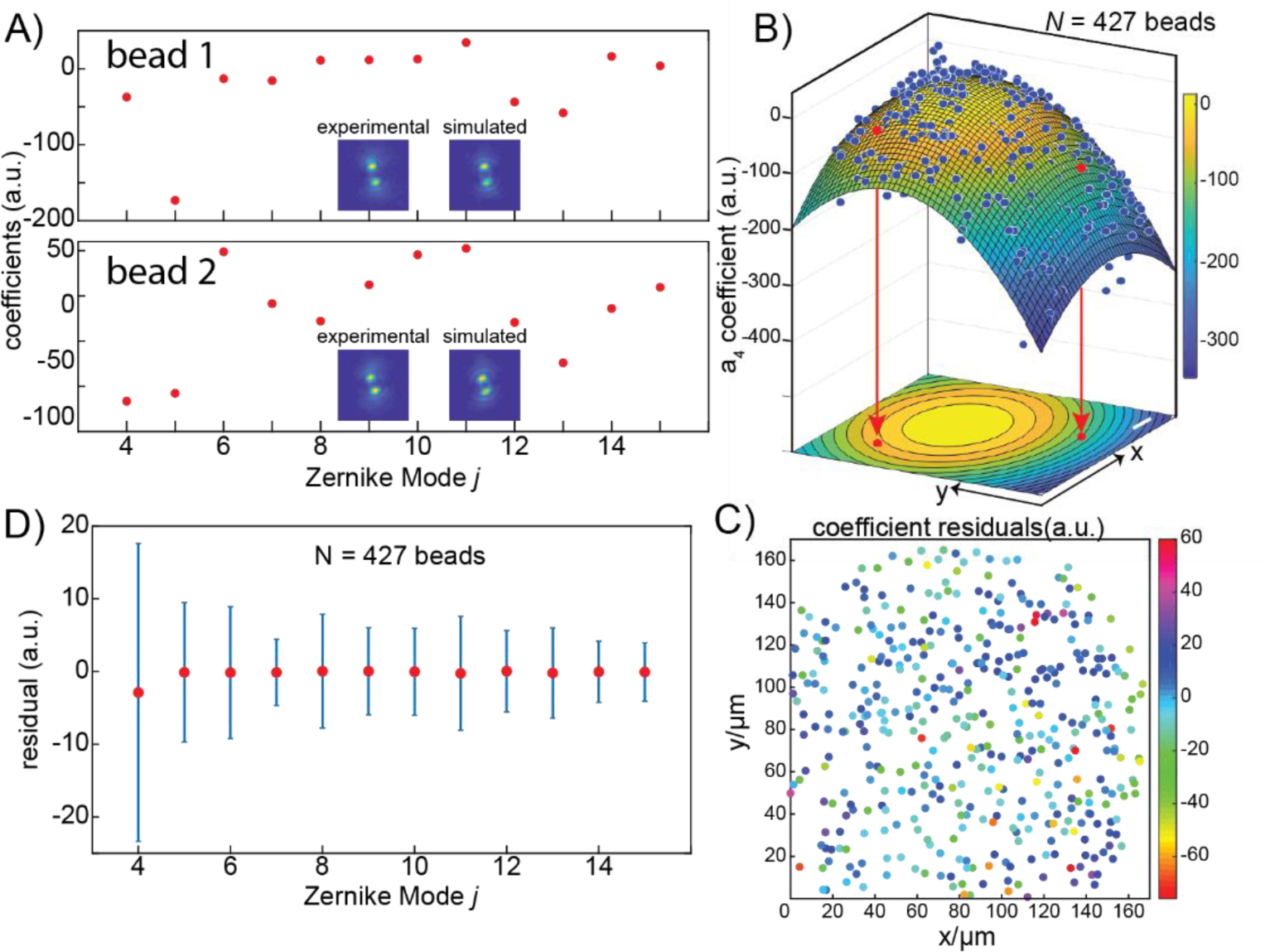
Spatially-variant Zernike coefficients. A) Coefficients for Zernike polynomials *j* = 4,…, 15 at two different bead positions in the field-of-view (indicated by red arrows and the red dots in panel B). Insets: experimentally measured DHPSFs (left) and simulated DHPSFs (right) based on the estimated Zernike coefficients. B) Spatial variation of Zernike coefficients a_4_ (defocus). The amount of defocus aberration were estimated at *N* = 427 bead positions (blue dots and red dots) pooled from three separately acquired fields-of-view. The surface is a second-order polynomial fit to the experimentally determined defocus coefficients. The corresponding contour map is also shown. Scale bar: 20 µm. C) The coefficient residuals, defined as the vertical difference between the blue dots and the polynomial surface. No spatial dependence is evident. D) Mean (red circles) and standard deviation (error bars) of coefficient residuals for all Zernike modes considered.

To verify the accuracy of the polynomial fits, we performed model cross-validation, compared estimates with different signal-to-background ratios, and performed simulations with beads at different *z*-positions. First, we randomly chose a subset of the acquired fields-of-view and performed the same polynomial surface fitting. The differences between surfaces created from all and subset of fields-of-view were not significant, i.e. they were within one standard deviations of the distribution of residuals [Fig. 3A]. Second, we performed the same estimation to the same field-of-view using experimental data with different signal and background levels. Again, we found that the resulting surfaces were the same within errors [Fig. 3B]. Third, we considered the effect of bead size heterogeneity on coefficients estimation via simulation. Aberrated images were simulated using the spatially-dependent aberration profile determined above and emitter *z*-positions of either 20 nm or 80 nm. The aberration coefficients were then estimated by assuming the *z*-position of 50 nm as described for experimental data. The difference of estimated coefficients relative to the ground truth was within one standard deviation of the coefficient residuals for every considered aberration except for defocus [Fig. 3C]. The defocus and primary spherical aberrations are strongly coupled with the distance *d* and emitter *z*-position, which explains the larger differences in the aberration coefficient *a*_4_ and *a*_11_. However, these simulations overestimate the bead size variation, which is 6 nm in diameter for the unstained microspheres according to manufacturer. Based on these experimental and simulated measurements, we conclude that polynomial surface fitting assuming a bead *z*-position of 50 nm is robust and yields accurate estimation of the spatially-variant aberration coefficient profiles through a large field-of-view.

**Fig. 3.**
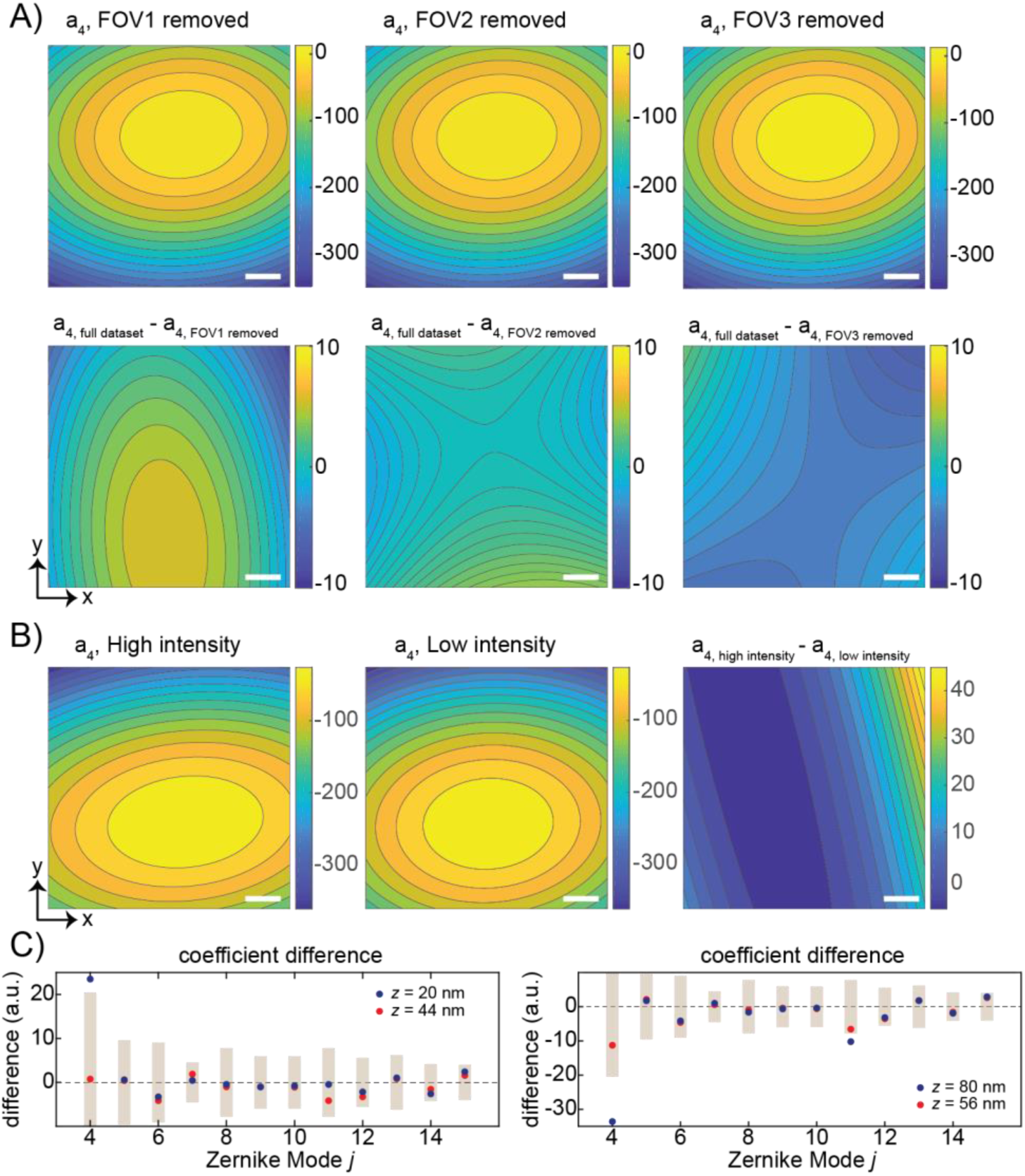
A) Cross-validation of polynomial surface fitting. The first row shows contour surfaces of the defocus aberration coefficient a_4_ obtained after removing the beads acquired in one of three fields-of-view. The second row shows the difference contours relative to the full dataset contour map shown in Figure 2B. Scale bars: 20 µm. B) Defocus aberration coefficient a_4_ contours in the other (green) channel of our microscope acquired using different laser intensities. For this dataset, the standard deviation of a_4_ residuals was 57 a.u. Scale bars: 20 µm. C) Effect of bead size on aberration coefficient estimation at a single position in the field-of-view. The difference between estimated and true coefficients are shown for different bead sizes (blue and red circles). Gray bars represent ± one standard deviation of the experimentally-determined coefficient residuals (Fig. 2D) centered at zero. Among the Zernike modes considered here, only the defocus (*j* = 4) and primary spherical aberration (*j* = 11) exhibit strong coupling with emitter *z*-position.

Finally, we performed the same estimation and fitting procedure in the second color channel of our microscope. The shapes of the corresponding surfaces between the two color channels are qualitatively similar, differing only in overall magnitude [Fig. 4]. We conclude that aberrations that are described by low order Zernike polynomials originate predominantly from optical components that are shared by both color channels of our microscope.

**Fig. 4.**
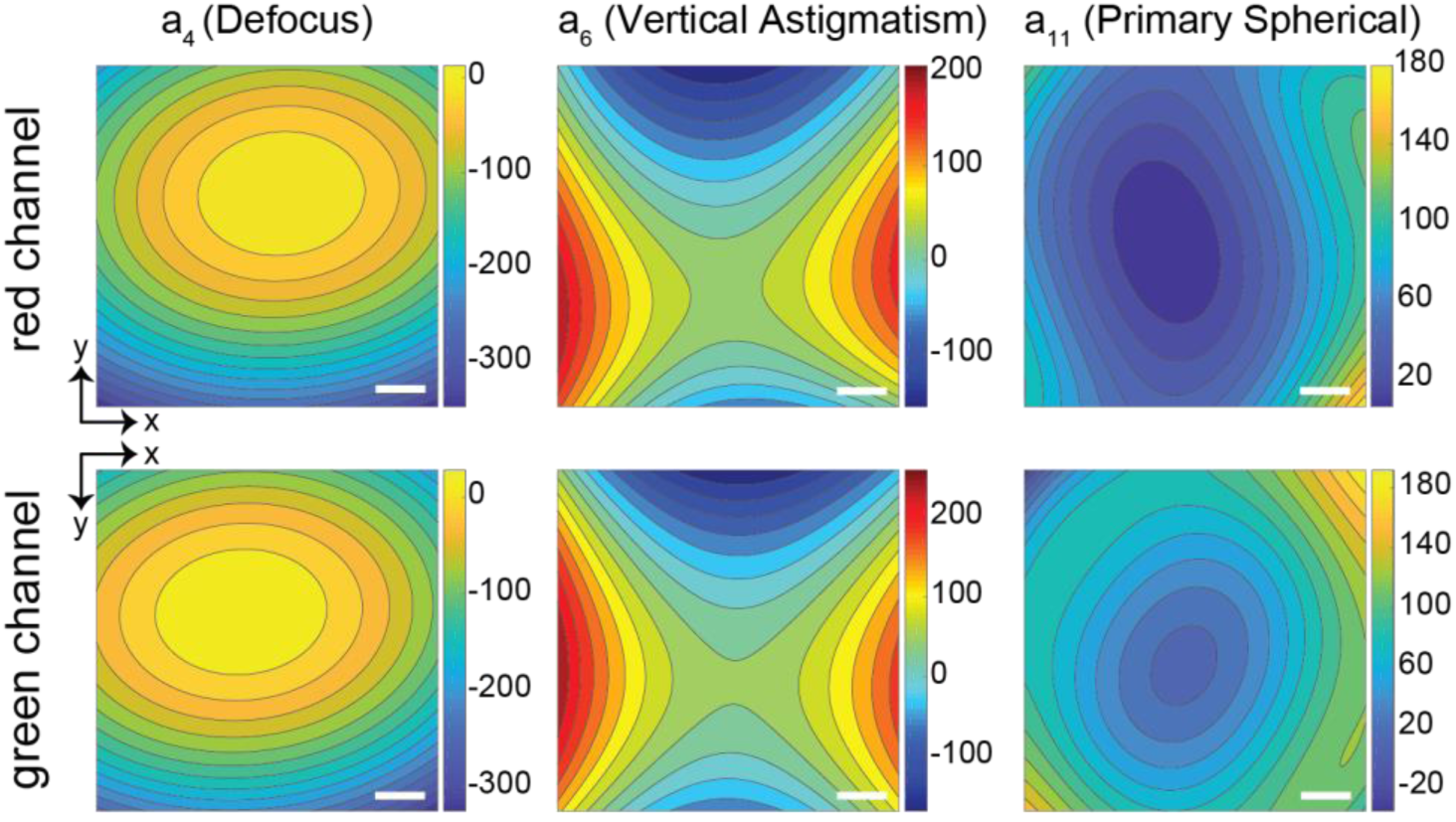
Aberration coefficient profiles in both color channels of our microscope. Top row: Aberration coefficient contour maps for a_4_ (defocus), a_6_ (vertical astigmatism) and a_11_ (primary spherical aberration) in the red channel. Bottom row: Corresponding contour maps in the green color channel. Scale bars: 20 µm.

### 3.2 Estimating the z-positions of Emitters on Flat Coverslips

To estimate the *x*,*y-*and *z*-positions of all beads in the field-of-view, the position of the nominal focal plane has to be determined in every frame. We estimated the distance *d* between the refractive index boundary and nominal focal plane using a reference marker with known *z*-position, such as a fluorescent bead of known size immobilized on the glass coverslip (*z* = 50 nm for a 100 nm fluorescent bead). Using the same reference marker in each frame, we then tracked changes in *d* over time due to stage drift [Fig. 5A]. Knowledge of the nominal focal plane position allowed us to estimate the *x*,*y-*and *z*-positions of all other beads in the field-of-view in every frame. We found that the estimated bead depths covered a wider range (stdev = 68 nm, range = [2, 343] nm) than expected, given the <10 nm bead size variation and the statistical localization precisions, defined as the standard deviations in the estimated *x*,*y-*and *z*-coordinates of each bead [blue localizations in top panel of Fig. 5B and Fig. 5C]. We noticed that several beads exhibited large localization precisions (colored in red in bottom panel of Fig. 5B and Fig. 5C). These beads were localized at larger depths and/or showed noticeable translational motion from frame to frame. Not considering these insufficiently immobilized beads, the averaged localization precisions were *σ*_*x*_ = 2.4 nm, *σ*_*y*_ = 2.9 nm, and *σ*_*z*_ = 3.6 nm.

**Fig. 5.**
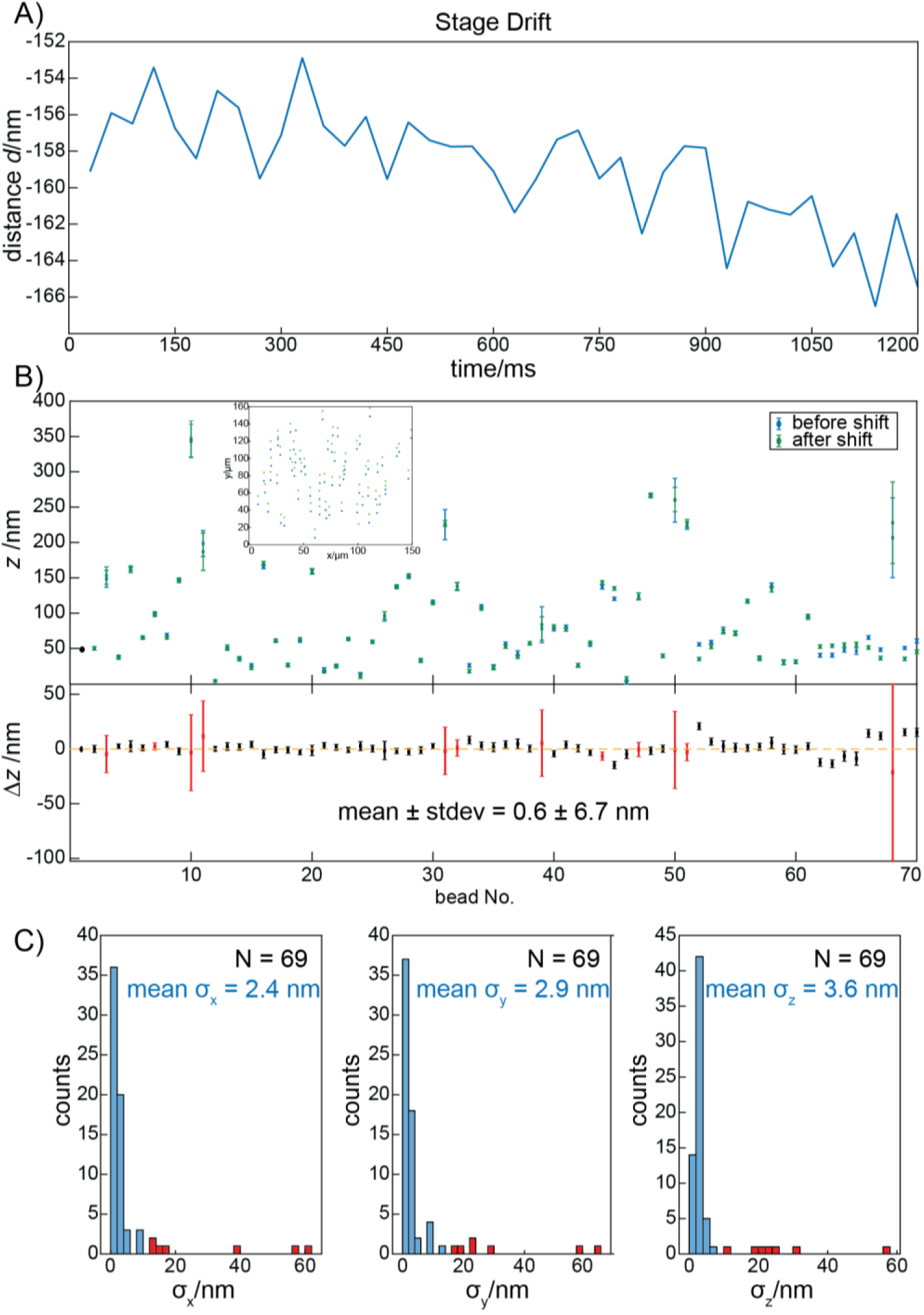
A) Change in the distance *d* due to stage drift. The same bead was used as a constant *z*-position reference. B) top panel: *z*-position measurements for different fluorescent beads on a flat coverslip immobilized by agarose before (blue) and after (green) a 10 µm lateral shift. The first bead (black) is the reference bead fixed at *z* = 50 nm. The red dots represent mean values from 40 measurement and the error bars represent the standard deviations. Inset: the spatial distribution of localized beads before and after the lateral shift. Bottom panel: the difference between *z*-position measurements before and after the lateral shift. Beads with noticeable motion from frame to frame in *x*,*y*- and/or *z*-directions are colored red. C) distribution of standard deviations of individual bead position measurement in the *x-*, *y*- and *z*-dimensions. Red colored bars represent beads with localization precisions of more than 10 nm (arbitrarily chosen threshold). The mean precison here is calculated excluding these aforementioned beads. If those beads were not excluded, the mean precision is: σ_x_ = 5.3 nm, σ_y_ = 5.7 nm, and σ_z_ = 5.1 nm.

The major contributor to the observed bead depth variance is the fact that some beads can get displaced into the agarose during sample preparation. We speculate that a second contributing factor is the thickness variations of the glass coverslip. Microscope coverslips are not polished to optical flatness, which for high quality optics can reach λ/20~25 nm. A third contribution could come from the use of inaccurate aberration coefficients. To rule out this possibility, we displaced the field-of-view by 10 µm and estimated the depths of the same beads again at their new positions in the field-of-view [green localizations in top panel of Fig. 5B,]. The average difference between bead depths before and after lateral displacement was 0.6 nm [Fig. 5B, bottom panel]. We conclude that our method is robust in estimating emitter depths accurately throughout the field of view. However, random thickness variations of the coverslip, which appear to be on the order of tens of nanometers, remain uncorrected.

### 3.3 Estimating the z-positions of Emitters in Agarose and Correlation between Two Color Channels

Imaging different components of the same structure in two different color channels provides information about their spatial proximity. However, due to different amount of aberrations present in different color channels, the images from two color channels are distorted differentially, limiting co-localization accuracy. Previous work has established the use of 3D transformation functions to correct the distortion and register two 3D super-resolution datasets acquired in different color channels [51]. In order to achieve 3D registration errors of less than 10 nm, transformation functions have to be computed based on large numbers of control points distributed throughout the 3D image volume. Such an approach leads to increased computational cost when imaging control points of similarly high density in an ultrawide field-of-view.

We hypothesized that the use of vectorial PSF models, with aberrations in two separate color channels calibrated, would yield depth accuracies sufficient to justify the use of 2D transformation functions to register 3D localization data. To test this hypothesis, we embedded fluorescent beads in agarose and compared their depths estimated in two color channels of our microscope [Fig. 6A, top panel, red and green plots, the same bead colored in black was used as reference marker]. The beads were distributed widely across the field-of-view [Fig. 6B], and the depths of the same beads matched up well with a mean difference of 0.8 nm and a standard deviation of 23.3 nm. We conclude that systematic *z*-colocalization errors can be eliminated and *z*-colocalization accuracies of about 20 nm can be achieved using the spatially-variant vectorial PSF model. We also note that the differences of more than 20 nm still occur for four of the beads considered here [bottom panel of Fig. 6A and Fig. 6C]. This observation may result from the existance of higher-order aberrations or sample-induced aberrations that are not accounted for by our model. To reduce *z*-colocalization errors into sub-10 nanometer regime, a locally calibrated 3D transformation function could be employed, possibly with lower control point densities [51], but we did not explore this option further.

**Fig. 6.**
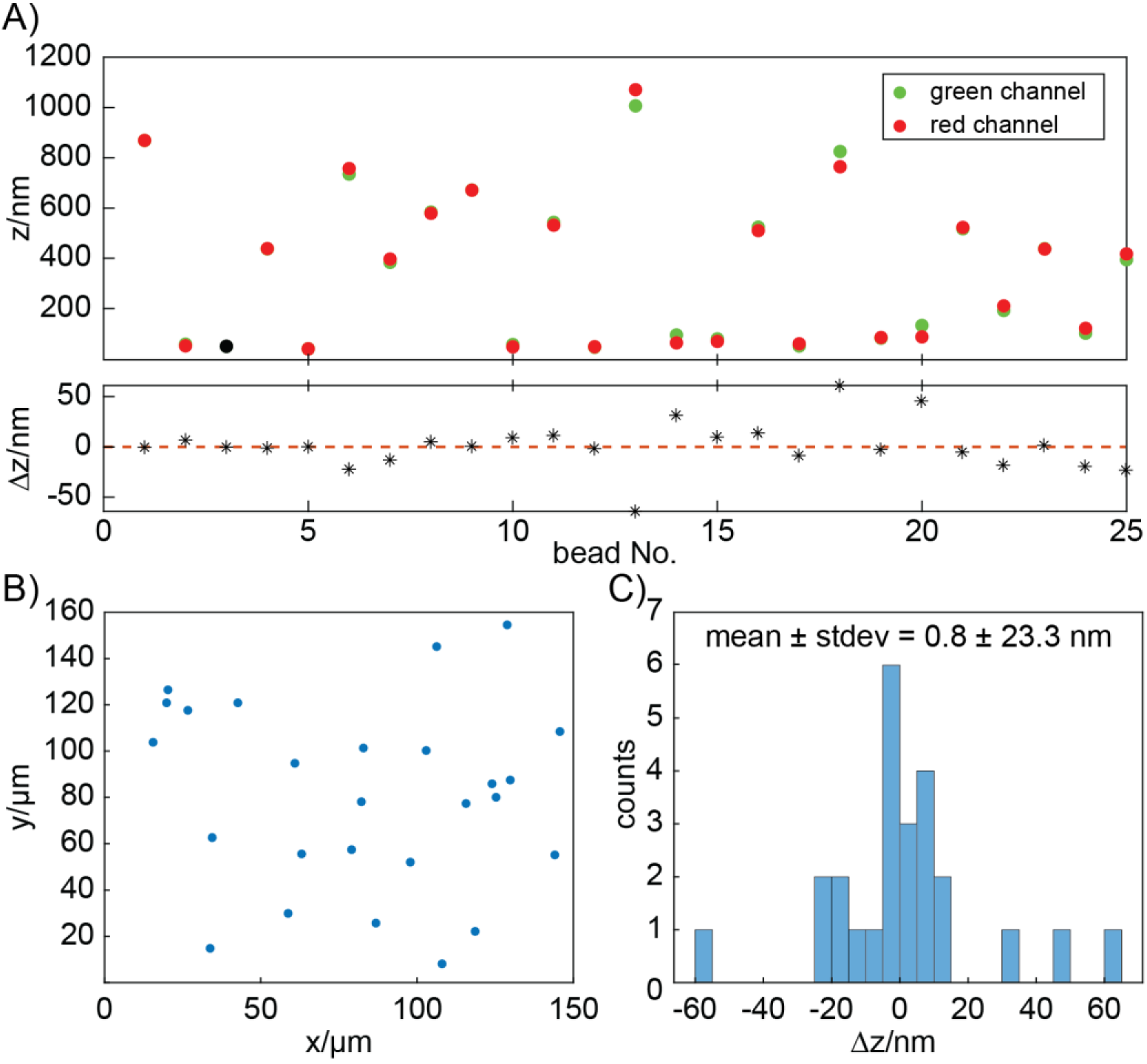
A) Top panel: *z*-positions of the same beads as measured in two separate color channels. The bead colored in black was used as a reference marker in both channels. Bottom panel: *z*-position differences for each bead shown in the top panel. B) The distribution of localized beads across the field-of-view. C) *z*-position differences.

### 3.4 Comparison to Simple Analytical Models

The above measurements suggest that localization precision of a few nanometers can be readily achieved at high signal-to-background ratios. To assess the improvements in accuracy and precision of position estimates made with the spatially-variant vectorial PSF model, we compared its performance to that of the double Gaussian model used in the Easy-DHPSF algorithm [53].

For experimental data of fluorescent beads immobilized on coverslips, we found that the Easy-DHPSF algorithm yielded bead *z*-positions that mapped out a bowl shape [Fig. 7A]. Emitters at the periphery were localized up to 800 nm higher than emitters near the center of the field-of-view because the spatially-variant aberrations, especially the defocus aberration, are not taken into account. We recapitulated this bowl shape using the Easy-DHPSF algorithm to analyze simulated images of bright emitters in the same *z*-plane imaged through an aberrated imaging system by using the experimentally-determined aberration coefficient profiles [Fig. 7B]. To simulate these images, isotropic emitters were placed adjacent to the refractive index boundary (*z* = 50 nm) and the nominal focal plane was made coincident with the refractive index boundary (*d* = 0). When using a global look-up table, spatially-variant aberrations are not taken into account and the *z*-error, defined as the average z-positions of *N* = 5 individual estimates relative to the ground truth, mapped out a bowl shape similar to the experimental measurement. When we refit the same data using many local look-up tables[37], the *z*-errors were reduced by roughly one order of magnitude [Fig. 7C]. However, this improvement was dependent on the emitter positions relative to the nominal focal plane. When the nominal focal plane was coincident with the refractive index boundary, most of the estimated z-errors were within the [-60, 80] nm range. By contrast, when the nominal focal plane was placed 500 nm above the coverslip (*d* = −500 nm), as would be done when localizing emitters inside a 1000 nm sized bacterial cell, the *z*-error range increased to [0, 200] nm [Fig. 7D]. Importantly, both the *d* = 0 nm and *d* = −500 nm cases, the *z*-error remains spatially-variant. By comparison, the spatially-variant vectorial PSF model estimates the *z*-positions without any spatial dependence (absolute mean z-error = 1.4 nm), because the *z*-errors are determined solely by the signal-to-background-limited localization precision. When the separation of the emitters and the nominal focal plane is random within a range of ±500 nm, the benefit of using many local lookup table is not as pronounced as for a constant separation distance [Fig. 7CD]. Such a situation is encountered when single-molecules randomly dispersed in a 1000 nm sized bacterial cells are localized relative to the nominal focal plane stably positioned at *d* = −500 nm [12]. The *z*-errors obtained by double Gaussian fitting using local look-up tables showed spatially dependent *z*-errors in the ranges of [0, 300] nm [Fig. 7E]. We found that the Easy-DHPSF algorithm often converges to incorrect estimates, especially when the simulated PSFs are severely aberrated. Incorrect estimates obtain frequently even when the double Gaussian model was heavily constrained and independent of whether global or local look-up tables were used. While grossly incorrect localizations, such as those outside the calibrated DHPSF *z*-range, can be filtered out during data post-processing, such filtering steps have the undesired outcome of reducing data acquisition throughput.

**Fig. 7.**
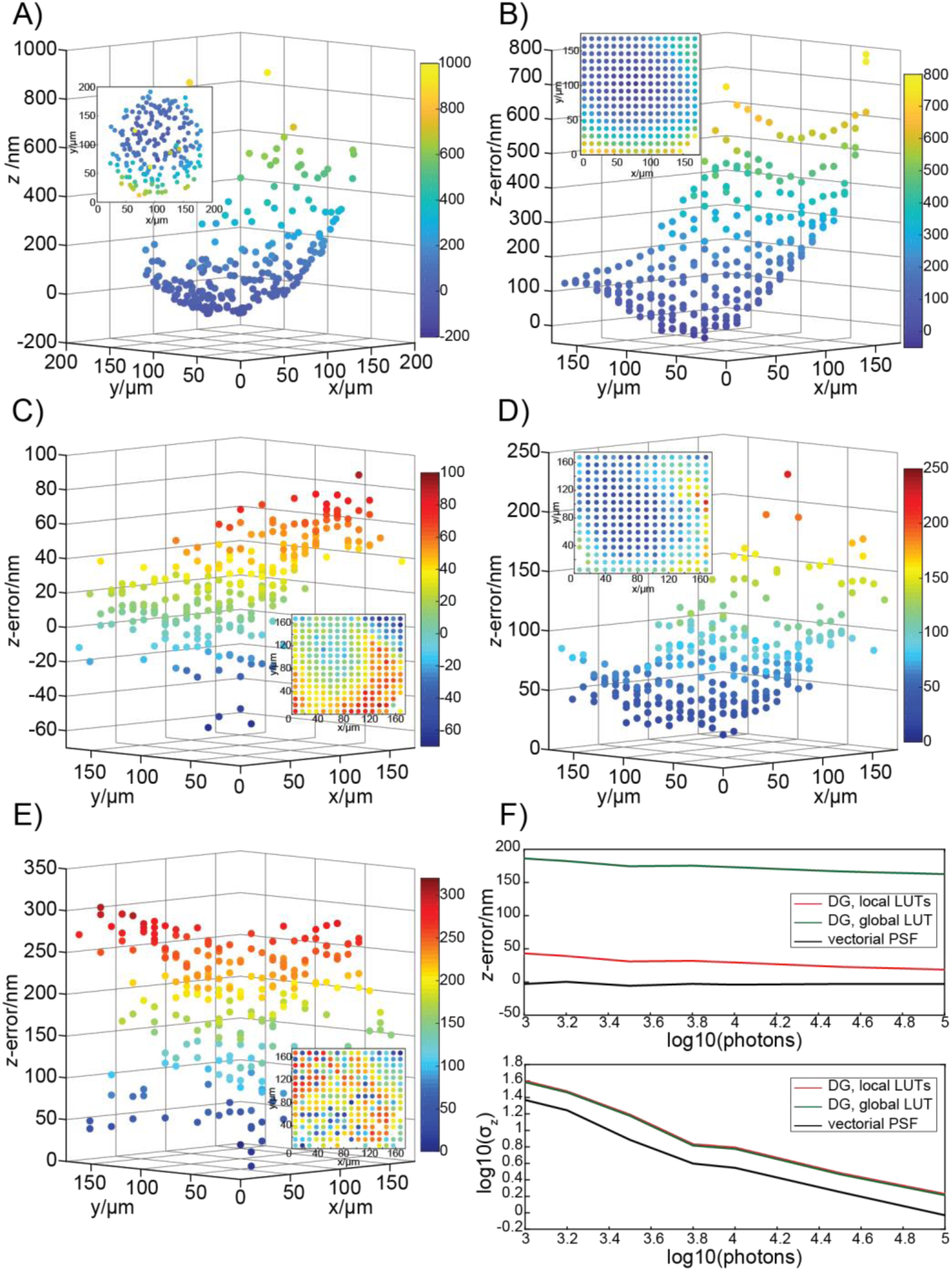
Comparison of double-Gaussian and the spatially-variant vectorial PSF model. A) *z*-positions estimates of fluorescent beads immobilized on the coverslip obtained by using the sdouble-Gaussian PSF model and a global lookup table. The *z*-positions map out a bowl shape with the beads near the center of the field-of-view localized with lower z-positions and the beads near the periphery localized with higher z-positions. Inset: Scatterplot of bead *x*, *y*-positions with each bead colored according to its *z*-position. B) Same results as in panel A, except that simulated DHPSFs of isotropic emitters were used as inputs. DHPSFs were generated on a 16-by-16 grid using the experimentally-determined aberration coefficients and assuming a constant emitter depth of 50 nm. The *z*-error, defined as the difference between estimated *z*-positions and the ground truth (50 nm), recapitulate the bowl shape observed experimentally. The nominal focal plane is coincident with the refractive index boundary (*d* = 0 nm). The range of z-error is consistent with the experimental result, i.e., [0 800] nm. Inset: the projection of emitters in *x*, *y*-plane with beads colored according to their z-errors. This applies to insets in panel C-E. C) The same simulation as in panel B, but the data were processed with a double Gaussian model using many local look-up tables. The *z*-error is reduced by roughly an order of magnitude, but still displays a spatial dependence. D) The same simulation as in panel C, but the nominal focal plane was placed 500 nm above the refractive index boundary (*d* = −500 nm). The range of *z*-error increased to [0, 200] nm, presumably due to increased separation between focal plane and emitters. E) The same simulation as in panel D (*d* = −500 nm, local lookup tables), but *z*-positions for DHPSF simulations were randomly chosen in the interval [0, 1000] nm. The range of *z*-error increased to [0 300] nm. F) Accuracy and precision comparison of double Gaussian vs. spatially-variant vectorial PSF model fitting. Noisy images of a single bead were simulated at different signal-to-background ratios at an off-center position in the field-of-view (70 µm lateral shift). The background was held constant at 10 photons per pixel. Both analytical models perform worse than the vectorial PSF model.

Finally, we evaluated the accuracy and precision of different methods as a function of the signal-to-background ratio. Noised images of a single bead (*z* = 50 nm; *d* = 0) were simulated with different signal-to-noise ratios and analyzed to extract the emitter *x*,*y-*and *z-*positions. At different signal-to-background ratios, the accuracy of the double Gaussian model can be improved by using a local look-up table, but the precison remains unchanged. For all signal-to-background ratios evaluated, the vectorial PSF model yields better accuracy and precision than the double Guassian model [Fig. 7F]. We conclude that the use spatially-variant PSF models is the optimal solution for single-molecule imaging in ultrawide fields-of-view, where optical aberration can vary substantially between different locations.

## 4. CONCLUSIONS

In this work, we introduce a spatially-variant vectorial PSF model to accurately and precisely determine the *x*, *y*- and *z*-positions of individual emitters in an ultrawide field-of-view. The spatially-variant aberrations are calibrated using bead images obtained at different positions in the field-of-view and at different nominal focal plane positions. This procedure quantifies the optical aberrations of the imaging system as linear combinations of Zernike polynomials. The coefficients of these linear combinations vary throughout the field-of-view and enable the generation of realistic PSF images for experimental data fitting.

We show that current implementation of the spatially-variant vectorial PSF model achieves better precision and accuracy than simple analytical models. Careful measurements in two color channels also showed that *z*-colocalization accuracies of 23.3 nm can be achieved. The spatially-variant vectorial PSF model presented here can be applied to model any (engineered) PSF corresponding to isotropic emitters, such as beads or rotating single-molecule fluorophores, or anisotropic emitters, such as fixed dipoles, in a refractive index mismatched medium. Limitations of the approach include the inability to calibrate the thickness variation of microscope coverslips *in situ*. Unknown undulations in the refractive index boundary limit the absolute *z*-localization accuracy to tens of nanometers.

Possible improvements of the vectorial PSF model could include incorporating additional Zernike modes beyond *j* = 15, and modeling refractive index variations in the specimen. Incorporating the finite bandwidth of the fluorescence emission spectrum into the PSF model may also help to further improve the agreement between simulated and experimental PSFs.

However, such improvements come at a rapidly increasing computational cost. Data processing time could be shortened through GPU computing as recently demonstrated for interpolated PSF models [45]. Alternatively, spatially-variant interpolated PSFs could be generated based on the method presented here. Such an approach would result in improved accuracy for estimating emitter positions, because locally-aberrated PSF models could be generated for interpolation by computationally moving the emitter to different *z*-positions instead of experimentally scanning the nominal focal plane through emitters at constant (and unknown) *z*-position.

